# Progesterone supplementation in mice leads to microbiome alterations and weight gain in a sex-specific manner

**DOI:** 10.1101/2021.10.06.463337

**Authors:** Meital Nuriel-Ohayon, Anna Belogovski, Sharon Komissarov, Meirav Ben Izhak, Oshrit Shtossel, Hadar Neuman, Oren Ziv, Sondra Turjeman, Shai Bel, Yoram Louzoun, Omry Koren

## Abstract

**Background:** Progesterone is a steroid hormone produced by the ovaries, involved in pregnancy progression and necessary for successful gestation. We have previously shown that progesterone affects gut microbiota composition and leads to increased relative abundance of *Bifidobacterium*.

**Results:** In non-pregnant female GF mice, levels of progesterone were significantly higher than in SPF mice of the same status. However, no significant differences were observed between GF and SPF males. Females treated with progesterone gained more weight than females treated with a placebo. In contrast to female mice, males treated with progesterone did not gain significantly more weight than males treated with a placebo. Progesterone supplementation led to microbial changes in females but not in males (16S rRNA sequencing). Accordingly, the weight gain observed in female mice treated with progesterone was fully transferable to both male and female germ-free mice via fecal transplantation.

**Conclusions:** We demonstrate that bacteria play a role in regulating progesterone levels in a female-specific manner. Furthermore, weight gain and metabolic changes associated with progesterone may be mediated by the gut microbiota.

## Background

The bi-directional interconnections between host hormones and the microbiome have been demonstrated in numerous studies^1-3^. While some hormones or hormone mimetics are produced by microbial species, hormones have been shown to reciprocally affect microbial populations, altering microbial composition and diversity. We have recently shown that progesterone modulates the pregnancy-associated gut microbial composition, in part, by increasing the relative abundance of *Bifidobacterium*^2^. This was seen in human cohorts, as well as female mice supplemented with progesterone, and following *in vitro* supplementation of fecal cultures with progesterone. The ability of progesterone to increase the growth of several bacterial species was previously demonstrated by additional *in vitro* studies, suggesting a direct effect of this hormone on the microbiota^2,4,5^. However, since elevation in progesterone levels occurs naturally in the context of pregnancy, it has been difficult to isolate and decipher the specific metabolic potential of the observed progesterone-related microbiome changes. Our current study focuses on a progesterone supplementation model in which other pregnancy-related factors such as hormonal, metabolic, or placental effects are eliminated. We hypothesized that progesterone-related microbiota alterations may lead to host metabolic changes including weight gain. This is based on the study by Koren et al (2012)^6^, demonstrating that fecal microbial transplants (FMT) from women during late pregnancy lead to increased weight gain in recipient germ-free (GF) mice. Associations between gut microbiota and weight gain are observed under additional situations including obesity, during early development, and following use of antipsychotic drugs or antibiotics^7-11^. Therefore, the progesterone-exposed gut microbiota may have a causative role in weight gain, as observed in pregnancy.

Another intriguing question regarding the effects of progesterone on the microbiome is whether these effects are sex specific. Here, we tested the microbial effects of progesterone supplementation in both male and female mice and the potential weight gain in both sexes following microbial transfer from progesterone-treated mice. Our findings demonstrate that progesterone significantly alters the gut microbial composition in females but not in males. Further, these progesterone-related microbial alterations lead to increased weight gain, which can be transferred to both male and female mice via FMT.

## Methods

### Experimental manipulations in mice

All experimental procedures were approved by the Bar-Ilan University Institutional Animal Care and Use Committee (protocol numbers 2015-11-29 and 20-04-2015). Mice (Swiss Webster) were bred and housed in either the germ-free or specific pathogen free (SPF) animal facilities at the Azrieli Faculty of Medicine, Bar-Ilan University, Safed, Israel. They were kept under a 12h light/dark cycle (light on 7:00, light off 19:00) and maintained at 22°C ±1. All mice were given free access to food and water and were fed from the same food batch (Harlan-Tekla, Madison, WI). Mice in the experiment were 8-10 weeks old. In each experiment, the day of randomization into groups was considered day 0.

### Progesterone implanted mouse model

SPF Swiss Webster mice were randomly assigned into one of two groups that were implanted with pellets either releasing (a) progesterone (n=9 males, n=9 females), or (b) placebo (n=9 males, n=9 females). They were housed 2-3 mice per cage. Pellets containing 35 mg progesterone or placebo (Innovative Research of America, Sarasota, FL) were implanted subcutaneously in the lateral side of the neck between the ear and the shoulder of the mice, and they released 1.67 mg/day for 21 days. Placebo pellets contained only a matrix without active progesterone. Throughout the experiment, weight and average food intake (3-4 mice per cage) were recorded every 3-4 days. Blood samples from the facial vein were collected on days 0, 11, and 21. Fecal samples from day 0 and 11 were taken for the FMT experiment described below.

### Fecal microbiome transplantation (FMT)

Fecal pellets from the progesterone implanted mouse study were kept at -80°C until use. Fecal pellets from donors (male and female mice receiving either a placebo or progesterone implant), were resuspended in 6 ml sterile PBS under anaerobic conditions, vortexed for 5 minutes, and allowed to settle by gravity for 5 minutes. Transplant into recipient germ-free Swiss Webster mice (8-10-week old, n=6 from each group) was performed by oral gavage using 200 µl of the supernatant. Pellets were not pooled; each recipient received FMT from a single donor, and the microbiota-recipient mice were housed separately under SPF conditions (1 mouse per cage) in autoclaved cages, with free access to autoclaved food and water. Fecal pellets and blood samples were taken on days 0, 7, and 14 for analysis of gut microbiota composition and blood progesterone levels, and the animals were weighed on the same days.

### Hormone analysis

Blood samples (100 μl) were collected from the facial vein using a sterile 5 mm goldenrod lancet (MEDIpoint, Mineola, NY) into collection tubes containing lithium heparin (Greiner Bio-One, Kremsmünster, Austria). Plasma was separated by centrifugation of blood samples at 1,500 × *g* for 15 min at 4°C and stored immediately at −80°C until analysis. Levels of progesterone in the blood were then measured with a chemiluminescent enzyme competitive immunoassay, using IMMULITE 2000 (Diagnostic Products Corporation, Los Angeles, CA).

### Fecal microbiota analysis

DNA was extracted from fecal samples using the PowerSoil DNA extraction kit (MoBio, Carlsbad, CA) according to the manufacturer’s directions following two minutes of bead beating. Purified DNA was used for PCR amplification of the variable V4 region (using 515F-806R barcoded primers) of the 16S rRNA gene, as previously described^2^. Amplicons were purified using AMPure magnetic beads (Beckman Coulter, Brea, CA), and subsequently quantified using the Picogreen dsDNA quantitation kit (Invitrogen, Carlsbad, CA). Equimolar amounts of DNA from individual samples were pooled and sequenced using the Illumina MiSeq platform at the Genomic Center of the Azrieli Faculty of Medicine, Bar-Ilan University, Safed, Israel.

### Bioinformatics and microbiome analysis

Microbial communities were analyzed using QIIME2 ^12^. Sequence reads were demultiplexed, and errors were corrected by DADA2^13^. Rarefaction was carried out using 10,900 sequences per sample.

### Normalization

Given the large variation in amplicon sequence variant (ASVs) values, we averaged all ASVs associated with the same taxonomy, and then transformed the resulting values by adding a minimal value to each feature (ASV) level (0.1) and calculating the 10-base log of each value. Statistical whitening was then performed on the table by subtracting the average and dividing by the standard deviation of each feature. The process was repeated for each mouse.

### Comparison between profiles

To calculate the average composition vector under different conditions, we used the average value of the abundance of each bacterium, following log normalization and Z scoring over the two experimental groups: PRO - progesterone treatment; PLC - placebo treatment. We also drew the vector representing the net effects, PRO - PLC derived by subtraction of FMT with placebo from FMT with progesterone treatment. This vector is the subtraction of two vectors, and as such represents the multiplicative (given the log normalization) net effect.

Finally, these (progesterone, placebo and the net effect) vectors were compared to a vector representing the relationship between bacteria and weight as represented by Weight_Corr: a result of Spearman correlation between bacterial composition and weight of the animals in the progesterone group.

We defined a direction for every profile based on the composition of bacteria. If two profiles were similar, they were aligned in the same direction. If the ratio between bacteria was similar in two profiles, but the variance was larger in one profile, then the profile with the larger variance was drawn longer. These profiles and the differences between profiles were plotted on the first two principal components.

### Statistical Analyses

Statistical analyses were performed using and PRISM 9.1.1. All data are expressed as mean ± SEM. False discovery rate (FDR) adjusted p-values were calculated for multiple comparisons.

## Results

### Lack of bacteria increases progesterone levels in females

As the relationship between microbiota and hormones is bidirectional, we first wanted to test whether the presence of bacteria influences progesterone levels in mice. To this end, we compared plasma progesterone levels of GF vs. specific pathogen-free (SPF) mice. We found that in non-pregnant female GF mice, the levels of progesterone were significantly higher than in SPF mice of the same status (mean ± SEM: 7,466±1,462 vs. 2,932.8±637 pg/ml, p=0.005). However, no significant differences were observed between GF and SPF males, which displayed similarly low levels of progesterone (2,846±1,214 vs. 2,487±1,466 pg/ml, p>0.05). These findings are intriguing, as they highlight the role of bacteria in regulating progesterone levels in a sex-specific manner (Fig. 1A).

**Figure 1.**
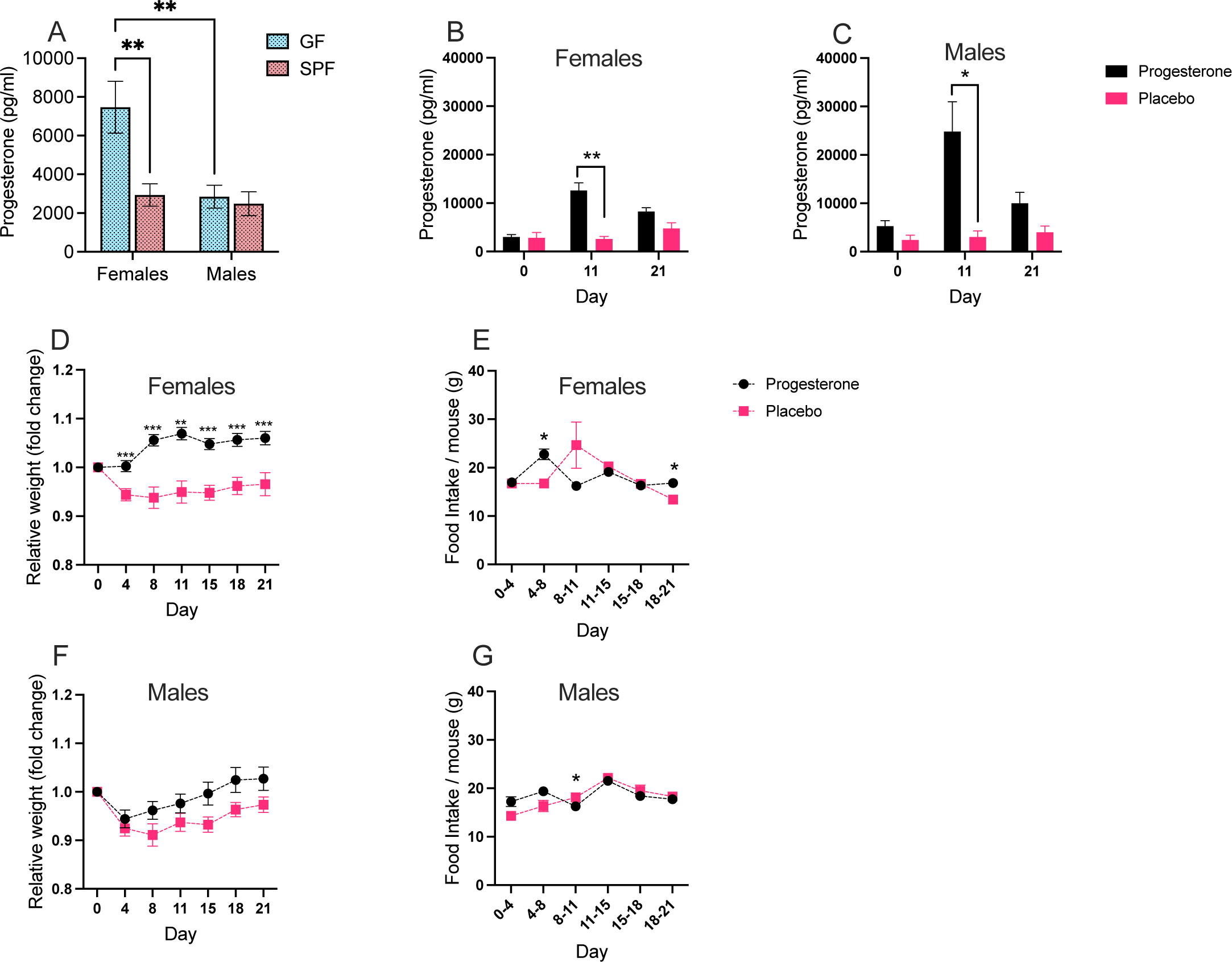
Supplementation of progesterone increases weight gain in female. (A) Progesterone levels in female and male GF (blue) and SPF (pink) mice. Progesterone levels in (B) female and (C) male mice after treatment with progesterone (black) and placebo (pink) pellets. (D) Weight gain and (E) food intake in females treated with progesterone (black) and placebo (pink) pellets. (F) Weight gain and (G) food intake in males treated with progesterone (black) and placebo (pink) pellets. Error bars indicate SEM; asterisks indicate statistically significant differences (*p<0.05, **p<0.001 ***p<0.001) as determined by the two-tailed student’s t-test.

### Progesterone increases weight gain in a sex-specific manner

To explore the potential effects of progesterone on gut microbial composition, we implanted progesterone (releasing ca. 1.67 mg/day) or placebo pellets subcutaneously for 21 days in both male and female SPF mice. In line with what we have previously shown^2^, blood progesterone levels were significantly higher both in female mice (Fig. 1B) that received progesterone compared to placebo and in males (Fig. 1C) 11 days after treatment.

In addition, female mice treated with progesterone gained more weight than female mice treated with placebo pellets. These changes were significantly different on days 4-21 (Fig. 1D). This was not due to increased food intake during the testing period (Fig. 1E); thus, progesterone treatment does not increase appetite, but rather, may increase the energy derived from the food. In contrast to the female mice, weight gain was not significantly different between male mice treated with progesterone and placebo (Fig. 1F). Similarly, no changes in food consumption were observed between the male mouse groups throughout the experiment (Fig. 1G).

### Progesterone leads to microbial changes in females but not in males

Using 16S rRNA gene sequencing, we analyzed the transplanted mice’s microbiome. We merged ASVs belonging to the same genus and log normalized the counts for each genus using the MIPMLP preprocessing tool^14^. To explore the global changes in the microbiome instead of specific bacterial shifts, we analyzed the projection of the microbiome on its main primary components following PCA. When projecting the log-normalized gut microbial composition on its first primary component and comparing progesterone vs. placebo-treated mice, we found noticeable differences between the female groups throughout the experiment, but not among the male groups (Fig. 1SA and 1SB, respectively) (p<0.05 for days 8, 21 for females).

We have previously shown that progesterone pellet implantation significantly changes the microbiome of female mice^2^. The results presented here indicate that the progesterone-related microbial alterations occur only in females, perhaps requiring additional female-specific components.

### Progesterone-supplemented microbiome transfers weight gain in both males and females

The above results reinforced our findings that progesterone supplementation in female mice leads to both a shift in microbial composition^2^, and to increased weight gain. We next wanted to confirm that microbial composition was the causal effect through which progesterone treatment contributed to weight gain. To this end, FMT was performed from female mice after 11 days of progesterone or placebo treatment, into male and female GF mice (Fig. 2A). Neither female nor male mice receiving FMT from females treated with progesterone showed elevated levels of progesterone themselves (Fig. 2B,C respectively). Interestingly, both female and male mice receiving an FMT from progesterone treated female mice gained significantly more weight, compared to their counterparts receiving FMT from placebo-treated mice 14 days following transplantation (Fig. 2D,E). However, fecal transplants from the same donor female mice before the intervention (at day 0), did not affect recipient weight (Fig. 2F,G), demonstrating that the supplementation of progesterone specifically induced a microbiome that promoted weight gain. This effect on weight gain was dependent on the sex of the donor; when feces from male mice treated with progesterone for 11 days was transplanted to both GF female and male mice, no changes in the weight of the recipients were observed (Fig. 2H,I respectively). Hence, these results reveal a female-specific effect of progesterone on microbiota composition, and this altered microbial composition can transfer weight gain to both sexes.

**Figure 2.**
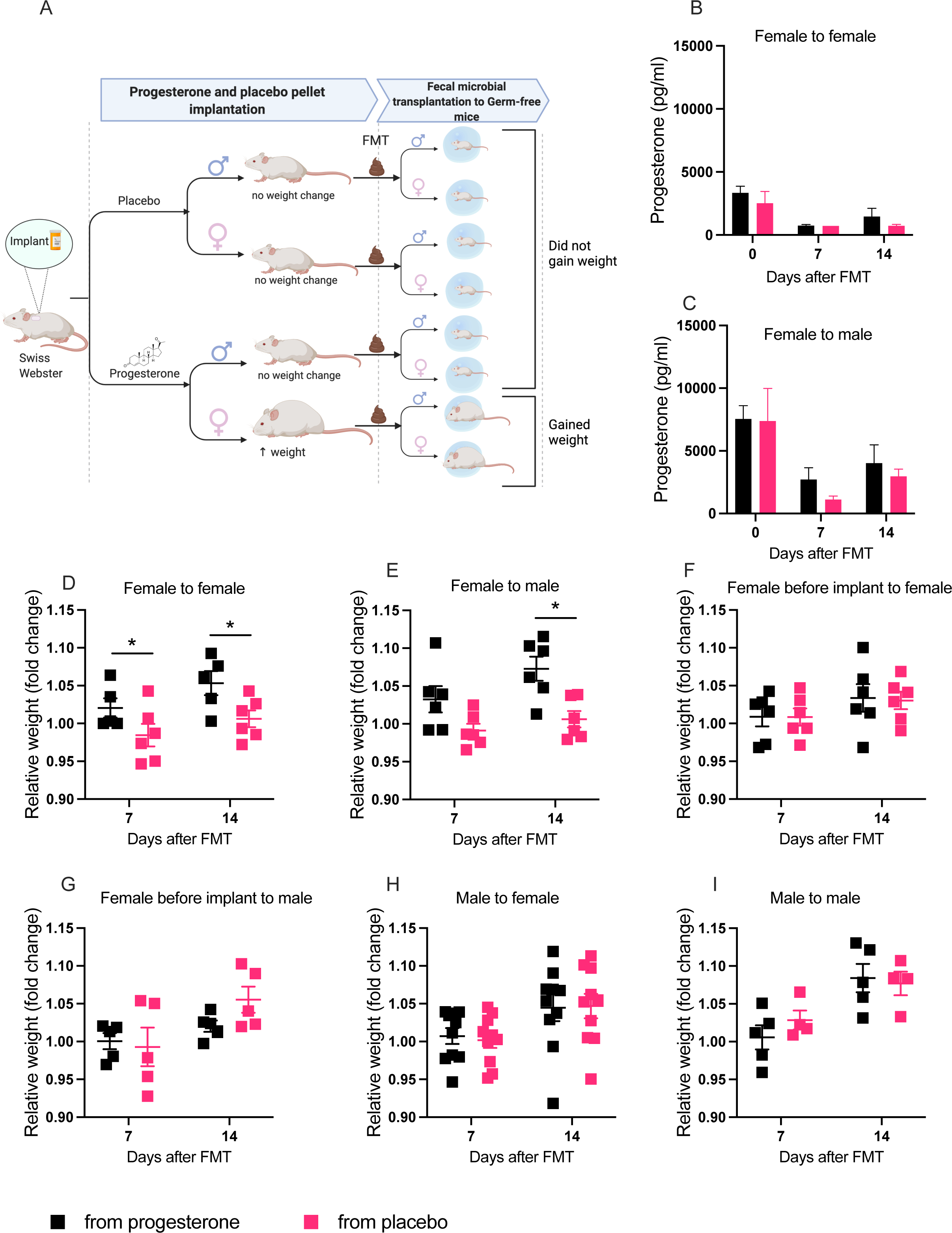
FMT from females treated with progesterone leads to weight gain in both male and female mice. (A) Experimental description. Progesterone levels in (B) females and (C) males after FMT from progesterone- (black) and placebo-treated (pink) female mice; relative weight in (D) females and (E) males after FMT from progesterone- and placebo-treated female mice (day 11). Relative weight in (F) females and (G) males after FMT from females before pellet treatment. Relative weight in (H) females and (I) males after FMT from progesterone- and placebo-treated male mice (day 11). Error bars indicate SEM; asterisks indicate statistically significant differences (*p<0.05) as determined by the two-tailed paired student’s t-test.

### Correlation of progesterone-altered microbiome and weight gain in mice

To relate weight change with the microbiome, we computed the Spearman correlation between the weight and each bacterial genus (normalized by frequency) in the placebo mice studied in the current experiment, producing a vector of correlations (Weight_Corr vector). To compare groups, we defined the average normalized weight profile (See Methods) for each group: PRO - progesterone transplants and PLC - placebo transplants. To separate the effect of the intervention itself from the placebo, we also computed the difference between PRO and PLC. Using this technique, the effects of progesterone on the gut microbiome composition, are separated from the effect of the transplant itself, as seen in the placebo, and possible bystander effects. The Weight_Corr and PRO - PLC vectors aligned for female but not male mice (Fig. 3A,B respectively). In female mice, the differences in microbial populations between the progesterone and the placebo groups (PRO - PLC) were highly similar to the vector representing the correlation between bacterial expression level and weight. In contrast, the placebo effect by itself (PLC) was almost opposite to the weight change vector. This implies a similarity between progesterone-related microbial alterations and the ones associated with weight gain. In male mice, there is no association between the weight change and transplant associated changes in microbiome.

**Figure 3.**
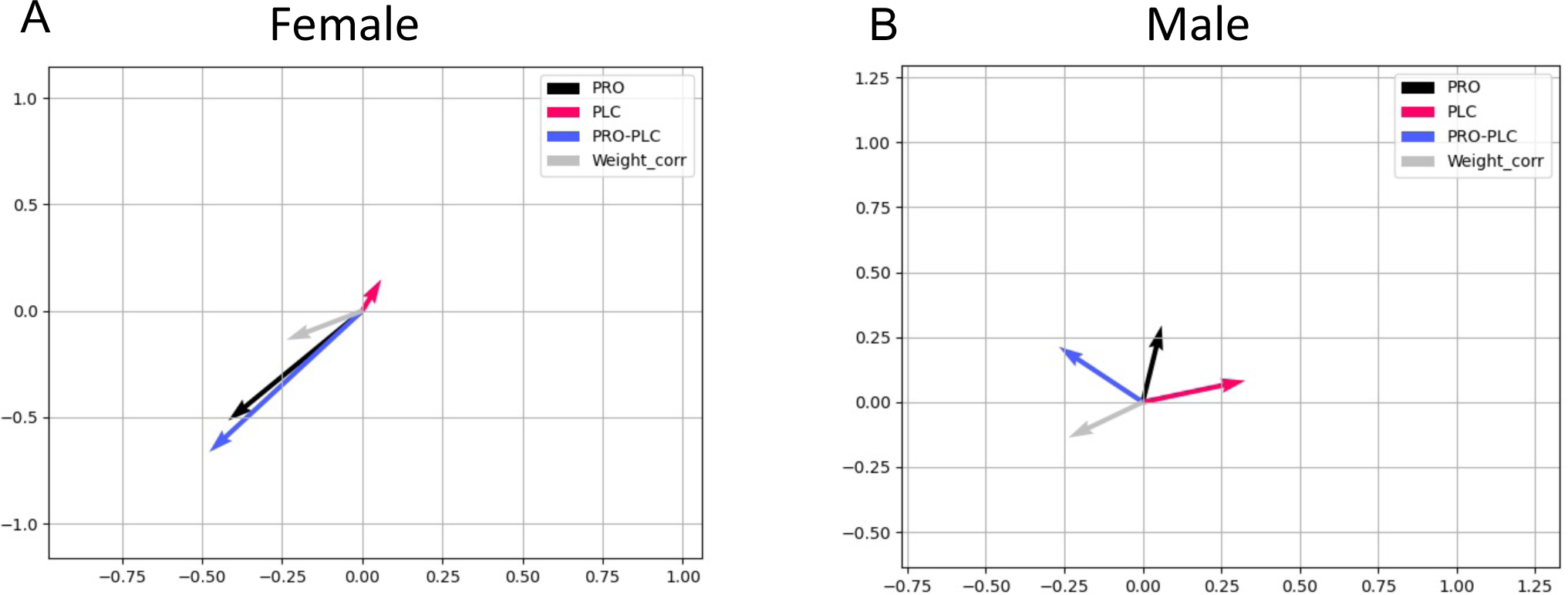
Supplementation of progesterone leads to microbial changes correlated with weight gain in female but not male mice. The projection on PC1 and PC2 of progesterone (PRO, black) and placebo (PLC, pink) treatments, the difference between them (PRO-PLC, blue), and weight correlation (Weight_corr, grey) vectors in (A) females and (B) males. It is clear that the impact of progesterone transplantation on females and males is different. While there is a significant effect of progesterone in females (represented by the directions of the pink and blue vectors), no relationships are observed in males.

## Discussion

The finding of higher levels of progesterone in GF compared to SPF female mice is intriguing as it suggests that bacteria play a role in regulating progesterone levels in SPF female mice and maintaining low levels in the non-pregnant state. The fact that progesterone levels are not elevated in male GF relative to SPF mice indicates a female-specific role of bacteria in regulating progesterone levels. One potential possibility is that in females, the microbiome interacts with a female-specific gene expression program to suppress progesterone levels. This could involve the expression of progesterone receptors, components in downstream signal transduction of progesterone, or additional female hormones.

Our findings show a significant increase in the body weight in female mice that received progesterone vs. placebo supplementation, as early as 4 days following onset of treatment. These findings are consistent with other studies showing that administration of progesterone to female mice results in weight gain^15^. A vast number of studies have shown that certain microbial compositions can themselves lead to increased weight gain in the host^11,16,17^. While several mechanisms may explain the effect of progesterone on weight gain, our results demonstrating weight gain in mice that received FMT from progesterone-vs. placebo-treated female mice strongly support direct involvement of the microbiota in this process. The specific mechanisms by which the gut microbiota may lead to weight gain are diverse and include involvement of the microbiota in the development of metabolic syndrome^18^, an effect on metabolism and immune functions, and the regulation of fat storage^19^. Another potential mechanism underlying gut microbiota-induced obesity is increased energy harvesting from food, which leads to an imbalance between energy intake and expenditure^20^. Gut microbiota can also increase the production of short-chain fatty acids, which are converted to lipids in the liver^11^. Additionally, higher levels of lipopolysaccharides (LPS), the major glycolipid component of the outer membrane of gram-negative bacteria, have been associated with increased weight; accordingly, adding LPS to a normal diet, induced insulin resistance and led to weight gain in mice^21^.

Interestingly, the influence of progesterone-exposed microbiota on weight gain was seen in both male and female mice throughout the FMT experiments. This emphasizes that while the effect of progesterone on microbial alterations is female-specific, the downstream effects of these microbial shifts on weight gain are not sex dependent.

To date, the influence of sex on microbiota composition is largely unknown. The changes are not dramatic, as most studies do not show clustering of microbiome composition according to sex. However, one study reported sex differences in human adults, showing larger colonization by *Bacteroides* in males than in females^22^.

The precise effect of progesterone on weight gain in males is unknown. Two studies previously reported a lack of effect of *in vivo* progesterone on male rat adipose tissue lipid metabolism^23,24^. This supports our findings and is apparently due to the absence of progesterone receptors in male rats. This is not surprising, as progesterone is a female hormone; and evidently, hormone receptor levels and metabolic mechanisms differ between males and females^25^. However, the changes in the female microbiota induced by progesterone exposure were enough to cause weight gain even in males, without the requirement for additional factors.

Pregnancy is a natural process in which progesterone levels rise dramatically to regulate and maintain gestation, for example, by preparing the uterine lining for implantation and preventing premature uterine contractions^26^. Progesterone is crucial for gestation, as mice lacking progesterone signaling are unable to establish pregnancy^27^. In parallel to the rise in progesterone, pregnancy is accompanied by weight gain and metabolic changes, as well as alterations in the oral, skin, vaginal, and gut microbial profiles, which partially resemble those observed in metabolic syndrome^28,29^. The co-occurrence of all these changes opens questions regarding their interconnections. Our results suggest that at least some of the microbial changes during late pregnancy occur due to the elevation of progesterone levels, and accordingly, at least some of the associated weight gain in pregnancy are likely be due to progesterone-driven bacterial alterations. Deciphering the reciprocal effects between microbiota and hormones is important and relevant not only for pregnancy but also for other conditions in which hormones are involved, such as progesterone supplementation as part of fertility treatments or hormone replacement therapy in menopausal women. These findings may yield a better understanding of mechanisms of weight gain following progesterone-mediated alteration of the microbiota and inform the design of specific probiotic treatments.

## Conclusions

Here, we found a counter effect in which bacteria play a role in regulating progesterone levels in a female-specific manner. Our findings also show a significant increase in the body weight of female mice that received progesterone vs. placebo supplementation, and this weight gain in female mice treated with progesterone was fully transferable to both male and female germ-free mice upon FMT. Altogether, our research suggests a bidirectional relationship between progesterone and the gut microbiota; progesterone levels are affected by gut microbiota on the one hand, while this hormone also affects gut microbiota composition on the other, further influencing host metabolism and weight gain.

## List of Abbreviations

ASV: amplicon sequence variant
FMT: fecal microbial transplantation
GF: germ-free
LPS: lipopolysaccharides
PCA: principal component analysis
PLC: placebo
PRO: progesterone
QIIME: quantitative insights into microbial ecology
SPF: specific pathogen-free
SW: Swiss Webster
rRNA: ribosomal ribonucleic acids

## Data availability

sequencing data is currently being submitted to a public repository.

## Declarations

### Ethics approval and consent to participate

All experimental procedures were approved by the Bar-Ilan University Institutional Animal Care and Use Committee (protocol numbers 2015-11-29 and 20-04-2015)

### Consent for publication

Not applicable

### Availability of data and material

sequencing data is currently being submitted to a public repository

### Authors’ contributions

MNO, OZ and OK designed the experiments; MNO and OZ performed the experiments; MNO, AB, SK, MBI, OS, HN, ST and YZ analyzed the data; MNO, HN, ST, SB, YL and OK wrote the manuscript; All authors reviewed the manuscript

## Acknowledgements

Not applicable

## Figure legends

**Figure S1**. Supplementation of progesterone leads to significant microbial changes in females but not in males. The difference in microbiome projection on PC1 (blue) over time in (A) females and (B) males after progesterone (black) and placebo (pink) supplementation.

