## Supplementary figures and images for "Progesterone supplementation in mice leads to microbiome alterations and weight gain in a sex-specific manner"

### Supplementary Figure 1

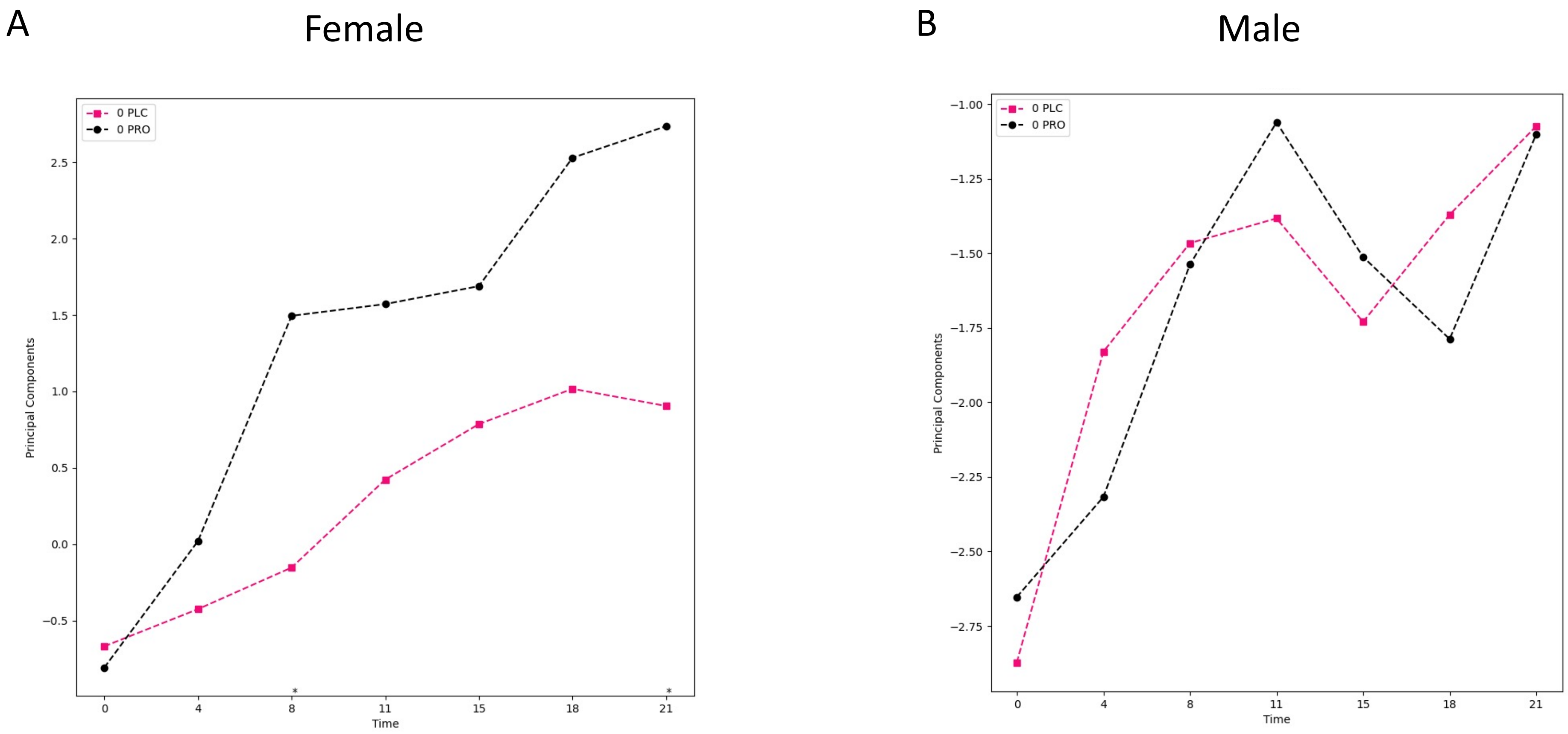
